# Growth Rate-Dependent Global Amplification of Gene Expression

**DOI:** 10.1101/044735

**Authors:** Rodoniki Athanasiadou, Benjamin Neymotin, Nathan Brandt, Darach Miller, Daniel Tranchina, David Gresham

**Affiliations:** Center for Genomics and Systems Biology; Department of Biology; Courant Institute of Mathematical Sciences, New York University

## Abstract

Regulation of cell growth rate is essential for maintaining cellular homeostasis and survival in diverse conditions. Changes in cell growth rate result in changes in rRNA and tRNA content, but the effect of cell growth rate on mRNA abundance is not known. We developed a new method for measuring absolute transcript abundances using RNA-seq, <u>SP</u>ike in-based <u>A</u>bsolute <u>R</u>NA <u>Q</u>uantification (SPARQ), that does not assume a constant transcriptome size and applied it to the model eukaryote, *Saccharomyces cerevisiae* (budding yeast), grown at different rates. We find that increases in cell growth rate result in increased absolute abundance of almost every transcript, with significant coordinated changes in abundances among functionally related transcripts. mRNA degradation and synthesis rates increase with increased growth rate, but to differing extents, resulting in the observed net increases in absolute abundance. We propose that regulation of ribosome abundance links environmental conditions to transcriptome amplification via nutrient-sensing pathways.

---

Control of gene expression by transcription factors, chromatin regulators^1^ and factors that act post-transcriptionally^2^ results in changes in mRNA abundance. Physiological properties of cells such as cellular growth rate^3-6^, size^7,8^ and ploidy^9^ have also been shown to affect gene expression programs. Typically, it is assumed that the majority of the transcriptome remains unchanged in response to different stimuli, and in different conditions or physiological states, and this assumption underlies standard normalization methods for comparative global studies of gene expression^10^. However, over-expression of the c-myc oncogene has recently been shown to result in globally elevated mRNA levels^11^ and variation of total RNA content has been previously described in mammalian cell cultures^12^. A global increase in total transcriptome pool size between samples violates assumptions underlying standard comparative differential gene expression analysis and requires the development of alternative approaches to global transcriptome analysis^13,14^.

Growth rate (GR) changes result in co-directional changes (i.e. a monotonic increase or decrease) in total cellular RNA content in both prokaryotic and eukaryotic cells. Previous studies found that changes in the total abundance of rRNA and tRNA contribute to the GR-dependent changes in RNA content in yeast cells^15,16^. Whether other types of RNAs, including mRNAs, are increased in absolute abundance with GR remains unknown. In this study, we aimed to determine the effect of GR on the absolute levels of all transcript classes and whether changes in rates of synthesis and/or RNA degradation underlie GR-associated changes in gene expression.

Rates of cell proliferation in unicellular and multicellular organisms vary dramatically between environmental conditions, cell types and developmental stages. Continuous culturing using chemostats^17^, in which a single essential nutrient is present at GR-limiting concentrations, provide an experimental means of maintaining exponentially growing populations at precise GRs. In a steady-state chemostat, the culture dilution rate is equal to the specific GR of the population (i.e. the exponential GR constant). We established steady-state cultures of *Saccharomyces cerevisiae* in carbon- and nitrogen-limited chemostats maintained at GRs of 0.12, 0.20, and 0.30 h^-1^ corresponding to population doubling times of 5.8, 3.5, and 2.3 hours respectively (**online methods** and Figure 1A). This experimental design enables discrimination of GR-specific from nutrient-specific effects on gene expression^4^.

**Figure 1.**
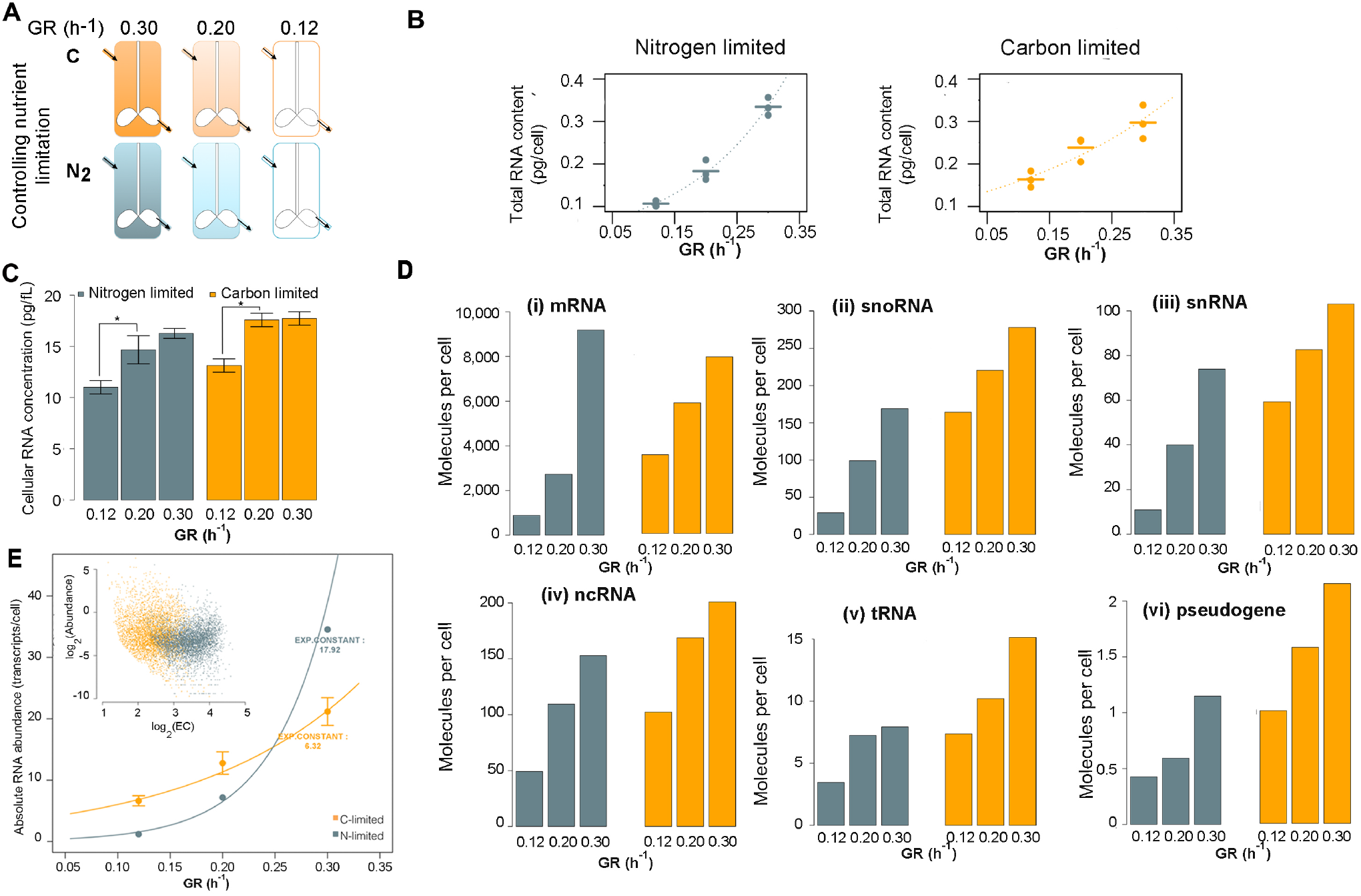
The total abundance of all RNA species increases with increased GR. A) Experimental design. Eighteen chemostats (three repeats of six experimental conditions) were established at the indicated GRs and nutrient limitations. The intensity of each chemostat color indicates the steady-state limiting nutrient concentration. B) Total RNA content per cell was estimated using quantitative extractions and performed in triplicate. Each biological replicate and their mean are shown. C) The concentration of RNA in the cell (fg/fL) increases with increased GR. The error bars represent the standard deviation of the mean across biological replicates. *p<0.05, t-test. D) The summed transcript abundance for each RNA category, measured by SPARQ. Samples grown in nitrogen limitation are blue and samples grown in carbon limitation are orange. In nitrogen and carbon limitation respectively: (i) 6408 and 6444 different detected transcripts (ii) 68 and 69 different detected transcripts (iii) 4 and 5 different detected transcripts (iv) 13 different detected transcripts each (v) 270 and 284 different detected transcripts (vi) 11 and 10 different detected transcripts. E) Use of SPARQ data for modeling the response to GR for an exemplary transcript, *ADE3*. Error bars depict SEM. The error bars for the nitrogen limited sample are too small to be drawn: SEM^GR=0.12^=4.51 10^−6^, SEM^GR=0.20^=1.16 10^−5^, SEM^GR=0.30^=2.33 10^−5^. Inset: the calculated exponential constants are independent of the transcript’s abundance (GR=0.12 h^-1^ plotted). F) GO enrichment analysis of the mRNAs with negative exponential constants. * adj.p<0.05, ** adj.p<0.01, *** adj.p<0.001, Bonferroni correction.

Consistent with earlier studies^15^, quantitative RNA extractions revealed a ~2.5-fold increase of total RNA (Figure 1B and **Figure S1A**) and rRNA (**Figure S1B**) content per cell as GR increases 2.5-fold. Although cell volume increases with GR too (**Figure S2**), we find increased total RNA concentrations in faster growing cells (Figure 1C, **Figure S1C** and **Figure S1D**).

To dissect effects of GR on the expression levels of all transcript classes we developed <u>SP</u>ike in-based <u>A</u>bsolute <u>R</u>NA <u>Q</u>uantification (SPARQ) (**online methods**). The method relies on adding a fixed amount of synthetic RNA spike-ins to a constant number of cells prior to RNA extraction (**Table S1, Figure S3**) followed by directional RNA-seq. The number of counts produced for each spike-in and its molar abundance (attomoles) informs a model relating the number of aligned reads for each gene to the number of transcripts per cell. By maximum-likelihood methods, we derive estimates of (a) the relative yield coefficient for each spike-in (number of aligned fragments produced per original RNA molecule) and (b) the attomoles of each RNA species multiplied by its relative yield coefficient. With the addition of a specific model for biological noise, we derive a complete statistical model for sequencing counts that allows parametric hypothesis testing and generation of synthetic data for model testing. Our normalization method is completely linear, unlike a recently published method^13^, and it does not seem to suffer from undue noise in the spike-in sequencing reads identified in a previous study^18^. Absolute mRNA abundances estimated using SPARQ are in good agreement with independent estimates from single-molecule FISH studies^19^ (**Figure S4**).

We find that the absolute abundance of all types of RNA, including mRNAs, increase with GR (Figure 1D and **Figure S5**) contrasting the results from a recent study^20^. In agreement with previous observations^21,22^ most mRNAs (97% - 64% of total detected) are estimated to be expressed at less than 1 molecule/cell (**Table S2** and **Table S3, Figure S6**). This suggests that only a fraction of cells express a given mRNA at any given moment in time, due to either cell cycle-related, or stochastic, gene expression.

Although tRNA transcripts comprise approximately 10% of the transcriptome^15^, our abundance estimates for this class of RNA are much lower (Figure 1D(v)). We attribute this to the presence of extensive post-transcriptional modifications in tRNAs that affect the efficiency of cDNA synthesis^23^, greatly reducing their sequencing yield relative to the synthetic spike-in RNAs and other endogenous RNAs. Despite lower sequencing yields an overall increase in the abundance of tRNAs is observed with increasing GR, consistent with earlier studies^15^.

To quantify the response of each mRNA to GR, we modeled absolute abundance as an exponential function of GR (**online methods**). The exponential constant (EC) from this model reflects the magnitude of absolute transcript abundance change with GR (Figure 1E) and is equivalent to the slope of a log-linear relationship between relative expression and GR used in a prior study^4^. We find that the majority of mRNAs respond significantly to GR (80 % in the nitrogen (**Table S2)** and 73% (**Table S3)** in carbon samples at 5% FDR). EC values are independent of absolute abundance and well correlated (r=0.53, p-value<2.2 10^−16^) between the two different nutrient limitations (**Figure S7B**) indicating that GR is the primary determinant of the response. Overall higher EC values in the nitrogen-limited samples indicate that the magnitude of the response to growth rate depends on the source of GR limitation (Figure 1E, inset).

Virtually all coding transcripts increase in absolute abundance with increased GR (**Figure S7A**). This predominantly positive response of mRNA abundance to GR contrasts with the study by Brauer *et al*^4^ in which the gene expression response to GR (quantified as a linear response of relative expression to growth rate) was centered around zero with an approximately equal number of genes responding positively and negatively to GR. This discrepancy between results is consistent with expectations when the assumption of a constant transcriptome size between conditions underlies normalization^13^. Indeed, inappropriate application of standard RNA-seq normalization and analysis to our dataset yields both positive and negative changes in gene expression with GR that are centered around zero (**online methods**).

Despite a concomitant increase in cell volume with GR (**Figure S2**), increasing absolute mRNA abundances result in increased cellular concentrations for the vast majority of mRNAs (**Figure S1D** and **Figure S8**), suggesting the GR effect on gene expression amplification is distinct from previously reported absolute mRNA abundance increases with increased cell volume^7^.

The broad range of calculated EC values (1.86-16.26 in carbon-limitation and 4.21-25.88 in nitrogen-limitation) indicates that absolute transcript abundance does not scale uniformly with GR, but that different genes respond to GR with different intensities. To assess the similarity of positive EC values (4,279 and 4,260 positive EC values, in nitrogen and carbon respectively) within sets of functionally related transcripts (*i.e*. the coherence) we used a modified gene set enrichment analysis algorithm (**online methods**, Figure 2). We found 61 gene sets in carbon-limitation and 107 gene sets in nitrogen-limitation that show significant coherence in EC values (**Table S4, Table S5**). These gene sets were manually grouped to seven broad categories as shown in Figure 2.

**Figure 2.**
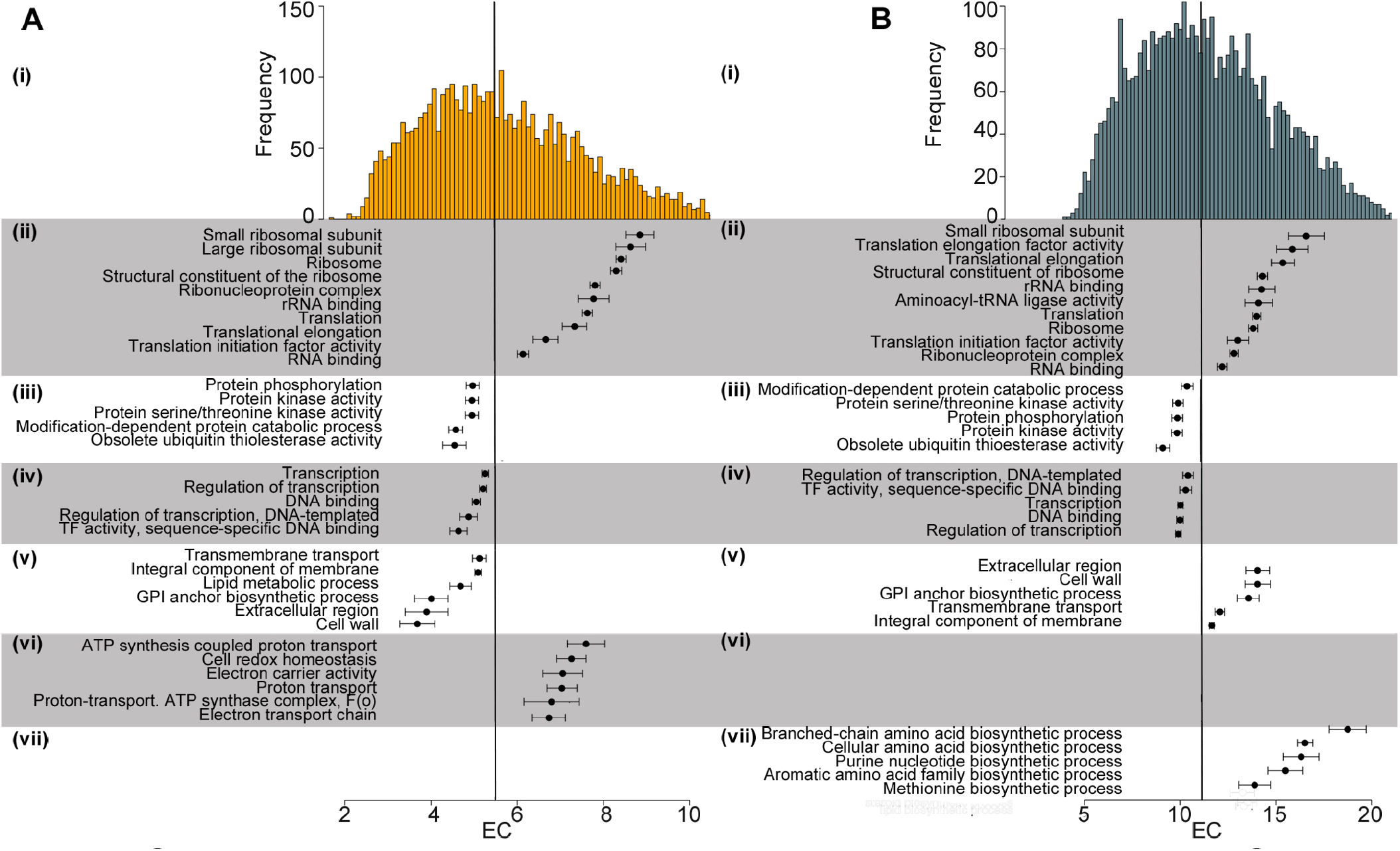
Gene set enrichment analysis of transcripts responding positively to GR. Distribution of positive exponential constants among mRNAs in **A**) carbon-limited and **B**) nitrogen-limited samples. The vertical line is the median response to GR. Panels (ii-ix): each point represents the average exponential constant of all genes in the gene set, error bars depict SEM. Gene Ontology super-groups are (ii) Protein synthesis, (iii) Post-translational modifications (iv) Transcription, (v) Cell membrane and cell wall, (vi) Proton-coupled energy production (vii) Amino acid biosynthetic process.

Genes involved in translation, especially those encoding ribosomal subunits, are highly coordinated in their response to GR and have high EC values (Figure 2(ii)). Additional groups of genes responding to GR in a coordinated way are related to post-translational modifications (Figure 2(iii)) and transcription (Figure 2(iv)). Gene sets associated with cell membrane and cell wall (Figure 2(v)) show unique behavior: although their response to GR is highly coordinated in both nutrient limitations, they are more sensitive to GR changes in nitrogen-limitation than carbon-limitation. This is consistent with our observation that cell size is much more sensitive to GR under nitrogen limitation (**Figure S2**).

The only cases in which we observed nutrient-specific coherence of EC values were genes involved in proton-coupled energy production (Figure 2(vi)) and biosynthesis of organic nitrogen biomolecules (Figure 2(vii)), under carbon- or nitrogen-limitation respectively. This likely reflects specific responses to the increased steady-state abundance of carbon and nitrogen with increased GR in carbon- and nitrogen-limited chemostats respectively.

Negative EC values reflect an inverse relationship between mRNA abundance and GR and are found for only a small number of transcripts (**Figure S7A)**. These transcripts show significant overlap between the two nutrient limitations (33 shared transcripts, p-value<10^−6^) and are enriched in genes involved in nutrient transport (**Figure S9**), which may reflect a common nutrient-scavenging response.

Changes in transcript abundance are the result of changes in the rate of RNA synthesis, RNA degradation, or a combination of both (modeled as *d[RNA]/dt = k -* ***α*** *\*[RNA]*, where k = synthesis, and ***α*** = degradation rate). We performed RATE-seq^24^ to determine the mRNA degradation rates at the two slowest GRs under nitrogen-limitation. We chose nitrogen-limitation because it elicits the strongest response to GR, which is most striking when comparing GRs of 0.12 h^-1^ and 0.2 h^-1^. We find that mRNA degradation rates increase for the vast majority of transcripts by an average of 3.1-fold. At steady state *(d[RNA]/dt = 0)* synthesis rates can be inferred from degradation rates and mRNA abundance *(k* = ***α*** *\*[RNA]*. Synthesis rates increase by an average of 7.7-fold, exceeding the increase in degradation rates and thereby accounting for the observed net increase in transcript abundance (Figure 3A).

**Figure 3.**
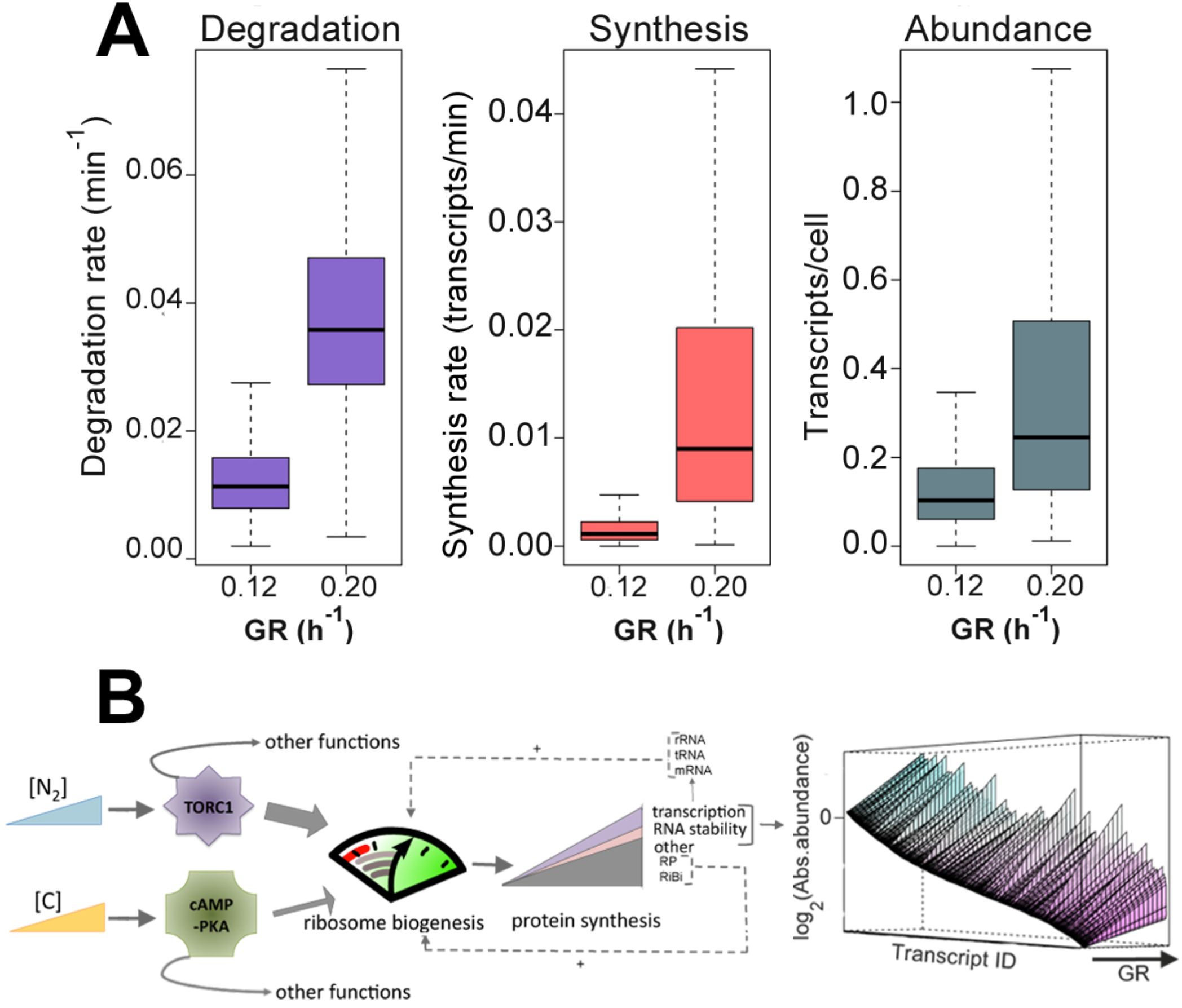
Kinetics of global gene expression amplification. **A**) RATE-seq was used to measure transcript degradation rates for all mRNAs at GRs of 0.12 h^-1^ and 0.20 h^-1^ in nitrogen-limited chemostats. Synthesis rates were calculated from the measured abundance and degradation rates. **B**) Proposed model of how regulation of ribosome biosynthesis results in globally increased RNA synthesis and degradation rates leading to global amplification of mRNA. Increasing concentrations of nitrogen and carbon activate the TORC1 and cAMP-PKA pathways respectively, which control ribosome biogenesis. A weaker effect of the cAMP-PKA pathway on activation of ribosome biogenesis (depicted by the different arrow weights), may explain the weaker response of gene expression to GR at the carbon-limited cultures. Increased ribosome biosynthesis and abundance leads to increased protein production rates including all proteins required for transcription and mRNA degradation. The two feed-forward loops created by the predicted accumulation of ribosomal proteins and RNAs may underlie the exponential increase of most mRNAs with GR. The net result of differentially increased RNA synthesis and degradation rates is a global increase in absolute mRNA expression levels.

Our results reveal a global effect of GR on absolute RNA levels produced by all three RNA polymerases that cannot be explained by a target-specific differential expression model. We propose that environmental control of ribosome abundance plays a central role in mediating transcriptional amplification and can explain our observations (Figure 3B). According to this model, elevated ribosome abundance results in increased translation rates and abundance of all proteins including RNA polymerases and RNA degradation components. Increases in these key enzymes lead to global increases in transcription and RNA degradation activity. Synthesis appears more responsive to GR than degradation, perhaps due to the protective effect of translation on mRNA stability^25^, leading to global transcriptome amplification.

Various lines of evidence support a universal role of regulated ribosome biosynthesis in determining global expression changes. Increasing nitrogen and carbon availability in our experimental conditions are known to activate the conserved TORC1 and cAMP-PKA pathways respectively^26^, both of which regulate GR and ribosome biogenesis. Ribosomes have been proposed to act as a mechanism coordinating GR and expression in bacteria^27^. In mammals, the c-Myc oncogene, over-expression of which leads to transcriptome amplification^11^, has been shown to also regulate ribosome biosynthesis^28^. Finally co-directional changes in mRNA synthesis and degradation have been observed under environmental stress^29,30^. Deciphering the effect of GR in baseline gene expression has important implications in understanding cell physiology and homeostasis, differentiation and dysregulation of gene expression. RNA-seq analysis using SPARQ provides a means of testing the extent of global gene expression amplification across a variety of conditions and cell types.

**Supplementary Figures** and **Methods** are available separately.

**Supplementary Tables S1-5** are available for download at: https://drive.google.com/folderview?id=0B231r3iakmyDSUZzWjFFZDREME0&usp=sharing

## Acknowledgements

We thank Jungeui Hong for assistance with the use of custom sequencing adaptors containing unique molecular identifiers (UMI). This work was supported by the National Institutes of Health (GM107466) and the National Science Foundation (MCB-1244219).

## Author contributions

R.A. conceived and designed the SPARQ assay, B.N. and N.B. prepared the samples and performed RATE-seq, B.N. and R.A. performed quantitative RNA extractions, R.A. performed SPARQ and data analyses, D.T. developed the SPARQ model, R.A., B.N. and D.G. were involved in the study design, D.M. performed mRNA-specific fluorescent labeling, R.A. and D.G. wrote the paper.

## Author information

The raw sequencing data used in this study are submitted to SRA with accession numbers SRP069877 (SPARQ) and SRP070446 (RATE-seq).

## References

1 Jaenisch, R. & Bird, A., Epigenetic regulation of gene expression: how the genome integrates intrinsic and environmental signals. Nat Genet 33 Suppl, 245–254 (2003).

2 Arraiano, C. M. et al., The critical role of RNA processing and degradation in the control of gene expression. FEMS Microbiol Rev 34 (5), 883–923 (2010).

3 Regenberg, B. et al., Growth-rate regulated genes have profound impact on interpretation of transcriptome profiling in Saccharomyces cerevisiae. Genome Biology 7 (11), R107 (2006).

4 Brauer, M. J. et al., Coordination of growth rate, cell cycle, stress response, and metabolic activity in yeast. Mol Biol Cell 19 (1), 352–367 (2008).

5 Castrillo, J. I. et al., Growth control of the eukaryote cell: a systems biology study in yeast. J Biol 6 (2), 4 (2007).

6 Ishii, N. et al., Multiple high-throughput analyses monitor the response of E. coli to perturbations. Science 316 (5824), 593–597 (2007).

7 Padovan-Merhar, O. et al., Single mammalian cells compensate for differences in cellular volume and DNA copy number through independent global transcriptional mechanisms. Mol Cell 58 (2), 339–352 (2015).

8 Wu, C.Y., Rolfe, P.A., Gifford, D.K., & Fink,, G.R., Control of transcription by cell size. PLoS Biol 8 (11), e1000523 (2010).

9 Galitski, T., Saldanha, A.J., Styles, C.A., Lander, E.S., & Fink,, G.R., Ploidy regulation of gene expression. Science 285 (5425), 251–254 (1999).

10 Dillies, M. A. et al., A comprehensive evaluation of normalization methods for Illumina high-throughput RNA sequencing data analysis. Brief Bioinform 14 (6), 671–683 (2012).

11 Lin, C. Y. et al., Transcriptional amplification in tumor cells with elevated c-Myc. Cell 151 (1), 56–67 (2012).

12 Darzynkiewicz, Z., Evenson, D.P., Staiano-Coico, L., Sharpless, T.K., & Melamed,, M.L., Correlation between cell cycle duration and RNA content. J Cell Physiol 100 (3), 425–438 (1979).

13 Loven, J. et al., Revisiting global gene expression analysis. Cell 151 (3), 476–482 (2012).

14 van de Peppel,, J. et al., Monitoring global messenger RNA changes in externally controlled microarray experiments. EMBO Rep 4 (4), 387–393 (2003).

15 Waldron, C. & Lacroute, F., Effect of growth rate on the amounts of ribosomal and transfer ribonucleic acids in yeast. J Bacteriol 122 (3), 855–865 (1975).

16 Kief, D. R. & Warner, J.R., Coordinate control of syntheses of ribosomal ribonucleic acid and ribosomal proteins during nutritional shift-up in Saccharomyces cerevisiae. Mol Cell Biol 1 (11), 1007–1015 (1981).

17 Ziv, N., Brandt, N.J., & Gresham, D., The use of chemostats in microbial systems biology. Journal of visual experimentation 80 (2013).

18 Risso, D., Ngai, J., Speed, T.P., & Dudoit, S., Normalization of RNA-seq data using factor analysis of control genes or samples. Nat Biotech 32 (9), 896–902 (2014).

19 Silverman, S. J. et al., Metabolic cycling in single yeast cells from unsynchronized steady- state populations limited on glucose or phosphate. Proceedings of the National Academy of Sciences 107 (15), 6946–6951 (2010).

20 Garcia-Martinez, J. et al., The cellular growth rate controls overall mRNA turnover, and modulates either transcription or degradation rates of particular gene regulons. Nucleic Acids Research (2016).

21 Marguerat, S. et al., Quantitative Analysis of Fission Yeast Transcriptomes and Proteomes in Proliferating and Quiescent Cells. Cell 151 (3), 671–683 (2012).

22 Holland, M.J., Transcript abundance in yeast varies over six orders of magnitude. J Biol Chem 277 (17), 14363–14366 (2002).

23 Zheng, G. et al., Efficient and quantitative high-throughput tRNA sequencing. Nat Meth 12 (9), 835–837 (2015).

24 Neymotin, B., Athanasiadou, R., & Gresham, D., Determination of in vivo RNA kinetics using RATE-seq. RNA (2014).

25 Huch, S. & Nissan, T., Interrelations between translation and general mRNA degradation in yeast. Wiley Interdiscip Rev RNA 5 (6), 747–763 (2014).

26 Broach, J.R., Nutritional control of growth and development in yeast. Genetics 192 (1), 73–105 (2012).

27 Scott, M., Gunderson, C.W., Mateescu, E.M., Zhang, Z., & Hwa, T., Interdependence of Cell Growth and Gene Expression: Origins and Consequences. Science 330 (6007), 1099–1102 (2010).

28 van Riggelen, J., Yetil, A., & Felsher,, D.W., MYC as a regulator of ribosome biogenesis and protein synthesis. Nat Rev Cancer 10 (4), 301–309 (2010).

29 Gasch, A.P., The environmental stress response: a common yeast response to diverse environmental stresses in *Yeast stress responses* (Springer, 2003), pp. 11–70.

30 Miller, C. et al., Dynamic transcriptome analysis measures rates of mRNA synthesis and decay in yeast. Mol Syst Biol 7, 458 (2011).

